# A behavioral polymorphism caused by a single gene inside a supergene

**DOI:** 10.1101/2020.01.13.897637

**Authors:** Jennifer R. Merritt, Kathleen E. Grogan, Wendy M. Zinzow-Kramer, Dan Sun, Eric A. Ortlund, Soojin V. Yi, Donna L. Maney

## Abstract

Behavioral evolution relies on genetic changes, yet few social behaviors can be traced to specific genetic sequences in vertebrates. Here, we show experimental evidence that differentiation of a single gene has contributed to divergent behavioral phenotypes in the white-throated sparrow, a common North American songbird. In this species, one of two alleles of *ESR1*, encoding estrogen receptor α (ERα), has been captured inside a differentiating supergene that segregates with an aggressive phenotype, such that *ESR1* expression predicts aggression. Here, we show that the aggressive phenotype associated with the supergene is prevented by *ESR1* knockdown in a single brain region. Next, we show that in a free-living population, aggression is predicted by allelic imbalance favoring the supergene allele. *Cis*-regulatory variation between the two alleles affects transcription factor binding sites, DNA methylation, and rates of transcription. This work provides a rare illustration of how genotypic divergence has led to behavioral phenotypic divergence in a vertebrate.

---

There is no doubt that many social behaviors are encoded in the genome. They are heritable, acted on by natural selection, and they evolve. Nevertheless, few genetic sequences have been directly linked to social behaviors in vertebrates. Most behavioral phenotypes are pleiotropic, and social behavior itself is flexibly expressed depending on context. This complexity, together with the many levels of biological organization separating a gene sequence from a social behavior, has made it difficult to completely understand why and how natural genotypic variation contributes to behavioral phenotypes.

The most promising animal models for identifying genetic targets of behavioral evolution are those with well-documented genetic variation linked to clear behavioral phenotypes. One such model is the white-throated sparrow (*Zonotrichia albicollis*), a North American songbird that occurs in two genetic morphs, white-striped (WS) and tan-striped (TS) (Fig. 1A). Whether a bird is WS or TS depends on the presence or absence of a naturally occurring series of rearrangements on chromosome 2, called ZAL2^m^. WS birds, which are heterozygous for ZAL2^m^, are more aggressive than TS birds, which are homozygous for the standard arrangement, ZAL2. ZAL2^m^ homozygotes, which are rare,^1,2^ are extremely aggressive.^2,3^ Thus, the ZAL2^m^ arrangement is strongly associated with aggression.

**Figure 1.**
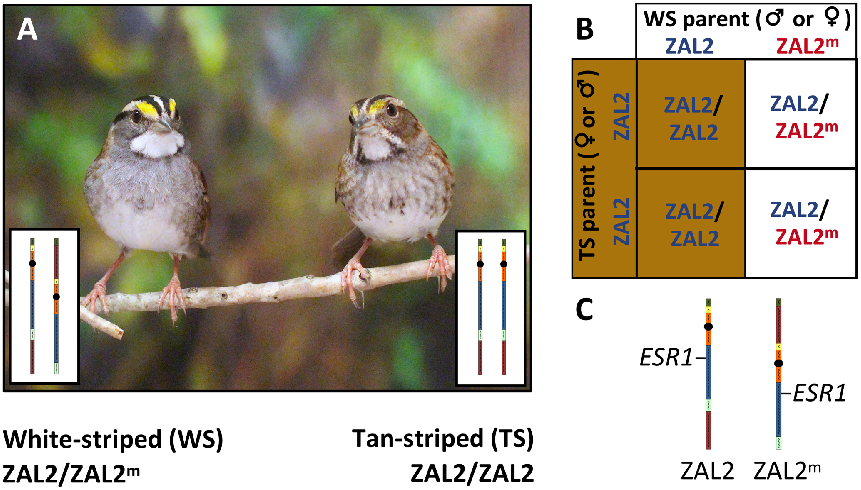
Polymorphism and disassortative mating in white-throated sparrows. (A) White-throated sparrows occur in two morphs: a more aggressive white-striped (WS) morph and a less aggressive tan-striped (TS) morph. WS birds are heterozygous for a rearrangement of chromosome 2, called ZAL2^m^, which contains a supergene. TS birds are homozygous for the standard ZAL2 arrangement. (B) Nearly all breeding pairs consist of one TS (ZAL2/ZAL2) bird and one WS (ZAL2/ZAL2^m^) bird. As a result, ~50% of the offspring are ZAL2/ZAL2^m^ heterozygotes and thus WS, and the rest are ZAL2/ZAL2 homozygotes and thus TS. (C) *ESR1*, the gene that encodes ERα, has been captured inside the supergene, resulting in the differentiation of the two alleles.^40^ Photo by Jennifer Merritt.

This species is unique among songbirds because of its disassortative mating system; nearly every breeding pair consists of one individual of each morph.^1,4,5^ Because almost all WS birds are heterozygous for ZAL2^m^, this mating system keeps the rearrangement in a nearconstant state of heterozygosity (Fig. 1B), profoundly suppressing recombination and driving genetic differentiation between ZAL2 and ZAL2^m^.^6^ The rearrangement is thus a “supergene” in that it contains a discrete set of co-inherited, co-evolving genes encoding a suite of traits that segregate together, in this case plumage and aggression.

One of the genes inside the rearrangement is *ESR1*, which encodes estrogen receptor alpha (ERα) (Fig. 1C). This gene is a particularly strong candidate for mediating the polymorphism in aggressive behavior. In sparrows, territorial aggression has been associated with plasma levels of testosterone,^7,8^ a steroid hormone that can be converted in the brain to the estrogen 17β-estradiol (E2); receptors such as ERα are therefore potential mediators of hormone-dependent aggression. Genetic differentiation between the ZAL2 and ZAL2^m^ alleles of *ESR1* has not caused non-synonymous substitutions that are likely to affect receptor function.^9^ Rather, it has introduced variation in regulatory regions that could alter levels of expression. The following observations suggest that variation in the expression of *ESR1* has functional relevance to the behavioral polymorphism. First, *ESR1* is differentially expressed between the morphs in several brain regions associated with social behavior.^9^ Second, expression in some of these regions predicts aggressive behavior better than does morph.^9^ Third, exogenous estradiol facilitates aggression in WS but not TS birds.^10,11^ Although these observations suggest that *ESR1* contributes to behavioral differences between the morphs, they provide only correlational evidence of such. To causally connect genotype to phenotype, we need evidence that morph differences in *ESR1* expression are (1) directly attributable to *cis*-regulatory variation in *ESR1* and (2) responsible for morph differences in behavior. The current set of studies accomplishes both of these goals.

## *ESR1* mediates an aggressive phenotype

In white-throated sparrows, *ESR1* expression differs by morph in several regions of the brain.^9,12,13^ The largest known morph difference is found in nucleus taeniae of the amygdala (TnA)^9,12^ (also called the ventrolateral arcopallium^14^), which shares molecular markers, connectivity, and function with the medial amygdala of mammals.^14,15^ In this region, *ESR1* expression is several fold higher in the more aggressive WS morph than in the TS morph. Further, this expression predicts aggression even when controlling for morph.^9^ We hypothesized that the morph difference in territorial aggression is caused, at least in part, by differential expression of *ESR1* in TnA. To test this hypothesis, we knocked down *ESR1* expression in this region in birds of both morphs. Local infusions of antisense oligonucleotides reduced the level of *ESR1* mRNA in WS birds to that of TS birds (Fig. 2A; Table S1,2). Whereas birds receiving scrambled control oligonucleotides showed the expected morph difference in *ESR1* expression (p = 0.009), there was no morph difference in birds receiving *ESR1* antisense oligonucleotides (p = 0.947). Treatment with antisense reduced levels of *ESR1* mRNA in WS (p = 0.048), but not TS (p = 0.994) birds (morph × oligo type, p = 0.041). Note that *ESR1* expression in TS birds is already low^9,12^, therefore knockdown can do little to decrease expression levels.

**Figure 2.**
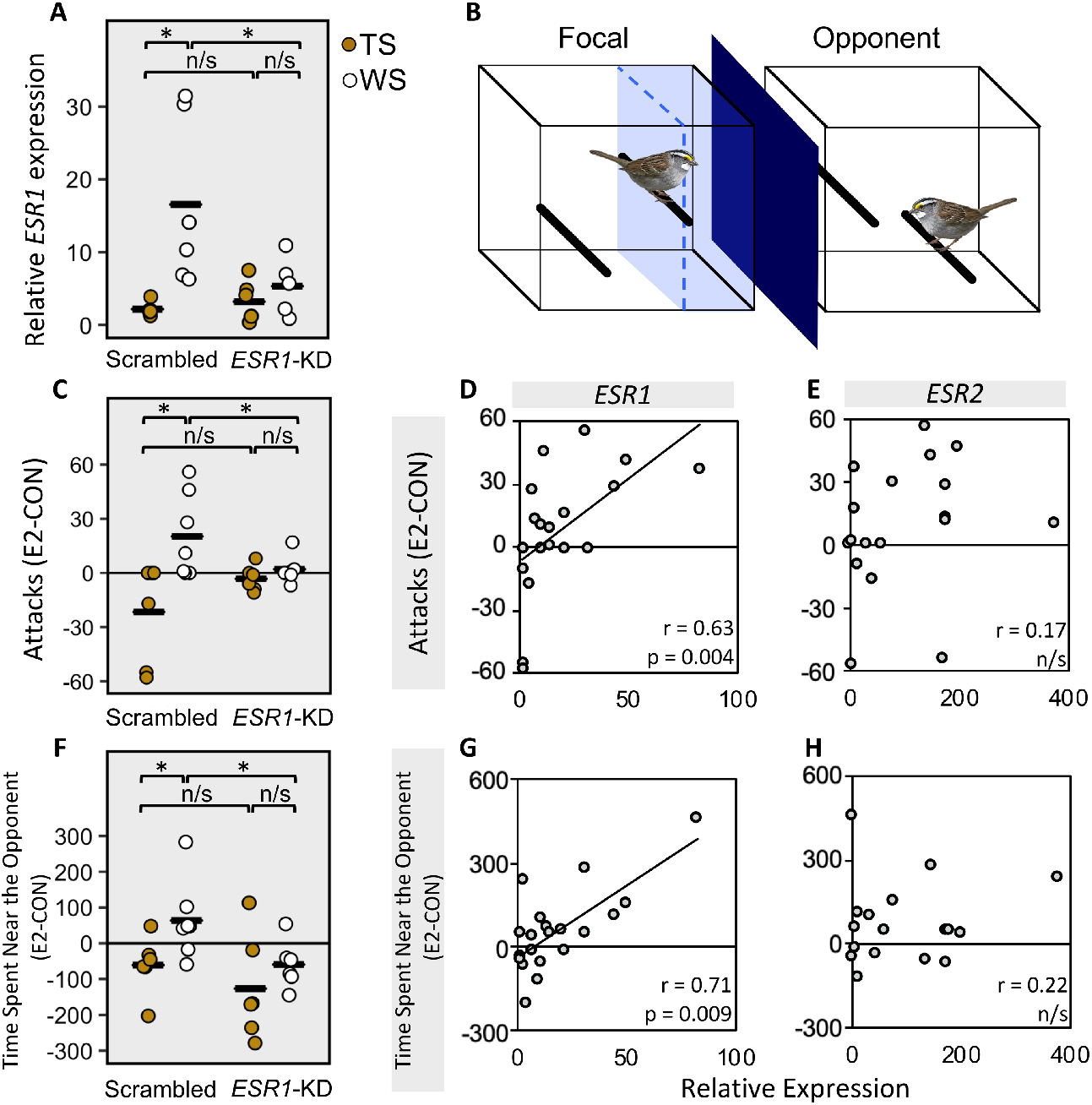
*ESR1* expression mediates the morph difference in aggression. (A) ESR1 expression in birds receiving infusions of scrambled control oligonucleotides or *ESR1* knockdown (*ESR1*-KD). Data are shown for pooled left and right TnA for each animal. (B) During behavioral testing, the cages of the focal bird and the opponent were separated by a visual barrier (dark blue; cage dimensions and bird not drawn to scale). Ten min after oral administration of 17β-estradiol (E2) or vehicle control (CON), the visual barrier was removed for 10 min. The Y-axis in C-H depicts the change in behavior between the CON and the E2 trials, in other words the degree to which E2 facilitated aggression in each animal. (C) *ESR1* knockdown significantly affected the degree to which E2 increased time spent near the opponent (defined as time spent in the area shaded in light blue in (B)) in WS birds only. This behavior was correlated with expression of *ESR1* (D) but not *ESR2* (E). (F) *ESR1* knockdown also significantly affected the number of attacks directed toward the opponent, again only in WS birds. This behavior was correlated with expression of *ESR1* (G) but not *ESR2* (H). Only birds that received infusions of scrambled oligonucleotides and the birds in which the cannulae missed TnA (n = 7) were included in the analysis of *ESR1* and *ESR2* expression. See Tables S1-4 for complete results. * p < 0.05.

Following knockdown treatment, we measured the extent to which treatment with exogenous E2 facilitated aggression towards another bird of the same species. Aggression was operationalized as time spent in proximity of a subordinate opponent and the number of attacks directed toward that opponent (Fig. 2B).^11^ The animals treated with scrambled, control oligonucleotides exhibited the expected morph difference,^11^ in that a bolus dose of E2 rapidly facilitated aggression in WS but not TS birds (morph × treatment interaction, p < 0.001 for both behaviors; Table S3,4). In contrast, knocking down *ESR1* blocked E2-facilitated aggression in WS birds (treatment × antisense interaction, p < 0.035 for both behaviors) (Fig. 2C,F). WS birds receiving knockdown exhibited levels of aggression indistinguishable from those of TS birds (p > 0.48 for both behaviors) (Fig. 2C,F). Thus, WS-typical levels of *ESR1* expression in TnA were necessary in order for E2 to facilitate aggression.

Whereas in our previous studies we measured aggression in free-living populations,^9,16,17^ in the present study we used in a lab-based assay.^11,18^ To confirm that *ESR1* expression in TnA explains aggression as operationalized in this lab-based assay, we tested for a correlation between aggressive behavior and the level of *ESR1* expression in TnA. Even when our analysis was limited to the control animals and “misses”, in other words animals not receiving knockdown in TnA, *ESR1* expression predicted aggression independently of morph (p < 0.01 for both behaviors) (Fig. 2D,G). We have thus shown that *ESR1* expression in TnA predicts a variety of aggressive behaviors in both field and lab.

The most highly expressed estrogen receptor in TnA is actually ERβ, not ERα.^17^ The gene encoding ERβ, *ESR2*, is not on chromosome 2 and its expression does not depend on morph.^17^ In this study, ERβ expression in TnA was unrelated to aggression (Fig. 2E,H), and ERα and ERβ expression in this region were not correlated (Fig. S1). Thus, despite its remarkably low expression in TnA, ERα seems to be the more important estrogen receptor driving the behavioral polymorphism.

## Allelic imbalance in ESR1 expression

We showed above that the aggressive phenotype of WS birds is mediated by their heightened expression of *ESR1*, compared with TS birds, in TnA. We hypothesized that the differential expression, which leads to differential behavior, is mediated by divergence of *cis*-regulatory regions of the *ESR1* gene. To test for differential regulation of *ESR1*, we quantified allelic imbalance (AI; Fig. S2), in other words the degree to which one allele is expressed more than the other, in TnA of free-living, behaviorally characterized WS birds (ZAL2^m^/ZAL2 heterozygotes). In the same birds, we looked for AI in two other regions of the brain: the rostral portion of the medial preoptic area (POM) and the hypothalamus (HYP, containing the caudal POM, paraventricular nucleus, anterior hypothalamus, and ventromedial hypothalamus). The expression of ERα differs between the morphs in these regions, albeit to a much lesser extent than in TnA.^9,12^

We detected AI in all three brain regions. The ZAL2 allele was overexpressed, relative to the ZAL2^m^ allele, in HYP and POM (Fig. 3A,D); in contrast, the ZAL2^m^ allele was more highly expressed than ZAL2 in TnA (Fig. 3G). Thus, among these three regions, AI favored ZAL2^m^ only in TnA. Although the degree of imbalance was not correlated with overall *ESR1* expression (quantified using qPCR) in HYP or POM, it was positively correlated with expression in TnA (Fig. S3). Our finding that the level of expression is predicted by the level of AI suggests that expression of the ZAL2^m^ allele, which is present only in the WS morph, is an important contributor to sensitivity to estrogens in this region.

**Figure 3.**
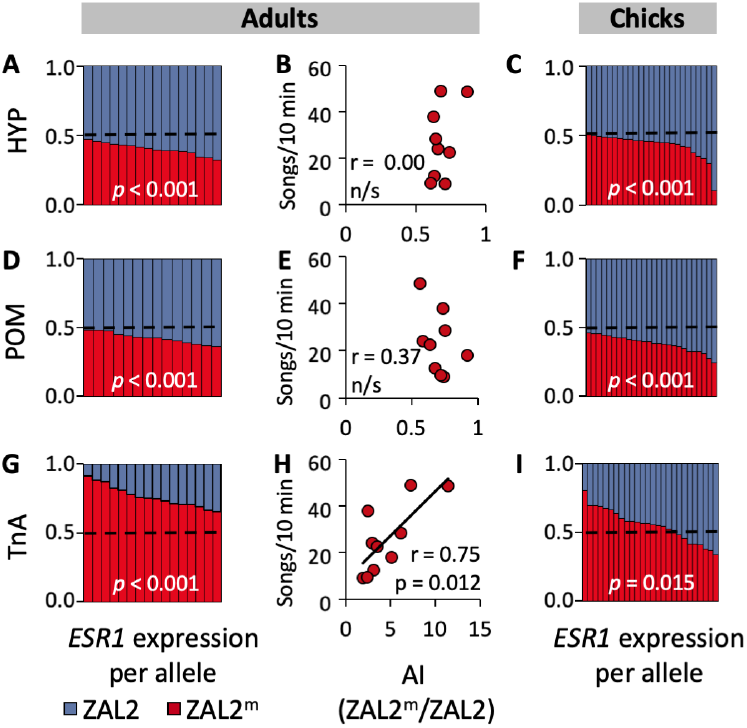
Allelic imbalance (AI) in *ESR1* expression in white-throated sparrows. We quantified AI in three brain regions in heterozygous (WS) adults (A,D,G) and nestlings (C,F,I). In the bar graphs, each column represents the relative percentage of ZAL2 (blue) and ZAL2^m^ (red) in a single bird. All regions in adults and nestlings showed significant AI (all p < 0.02; Tables S5,6). (B,E,H) Behavioral responses of adult males to simulated territorial intrusion were not predicted by the degree of AI in HYP (B) or POM (E); however, they were significantly correlated with AI in TnA (H). POM, medial preoptic area. HYP, hypothalamus. TnA, nucleus taeniae of the amygdala.

In free-living male white-throated sparrows, expression of *ESR1* in TnA predicts aggressive responses to simulated territorial intrusion (STI); in fact, this expression predicts the behavior even better than morph itself.^9^ In this study, we tested whether aggression in response to STI is predicted by the expression of the ZAL2^m^ allele specifically, relative to the ZAL2. We found that the degree of AI in TnA, which is ZAL2^m^-biased (Fig. 3G), strongly predicted aggressive responses to STI (Fig. 3H). In contrast, allelic imbalance in HYP and POM did not predict those responses (Fig. 3B,E). Thus, the degree to which a bird engaged in territorial aggression, which was markedly higher in the WS than TS birds,^9,11,16^ was predicted not only by overall *ESR1* expression in TnA but also specifically by the relative expression of the ZAL2^m^ allele, which is present only in WS birds.

Engaging in territorial aggression can affect plasma levels of sex steroids^19^ and presumably expression of steroid-related genes. Thus, is it possible that the correlation between aggressive behavior and *ESR1* expression is caused by the propensity of WS birds to engage in more territorial behavior. We therefore tested whether the pattern of AI in adulthood is already present early during development, before birds are engaging in territorial aggression. We quantified the ZAL2^m^/ZAL2 ratio in TnA, POM, and HYP in nestlings at posthatch day seven, two days prior to natural fledging. AI was detected in nestlings in all three regions (all p < 0.02; Fig. 3C,F,I; Table S5,6). As was the case for adults, the ZAL2^m^ allele was expressed more than ZAL2 in TnA (Fig. 3I), but in HYP and POM the ZAL2 was expressed more (Fig. 3C,F). Overall expression of *ESR1* in TnA is higher in WS than TS nestlings at the same age,^12^ thus both the allelic imbalance in WS birds and the morph difference in *ESR1* expression precede the behavioral differences in adulthood.

## Cis-regulation of ESR1 expression

Above, we showed that *ESR1* expression in TnA is causal for an aggressive phenotype in WS birds and that this aggressive phenotype is predicted by expression of the ZAL2^m^ allele specifically. We next explored the genetic and epigenetic mechanisms underlying AI. First, we hypothesized that *cis*-regulatory divergence has led to differential transcriptional activity of the two alleles, potentially because of divergence of transcription factor binding sites. Second, we hypothesized that epigenetic regulation of the *ESR1* promoter regions differs between the alleles, resulting in differential expression. We examined the regulatory regions upstream of transcribed portions of *ESR1* to identify variation likely to cause differential regulation of the alleles. *ESR1* contains three transcription start sites (TSSs) that drive the expression of transcripts that are expressed in the brain in both morphs (Fig. 4A).^9^ The transcript 1C (5’ UTR only) or the transcript from 1B (5’ UTR and exon 1B) can be spliced at +1 bp onto 1A (5’ UTR and exon 1A), generating three possible mRNA isoforms: 1C^1A, 1B^1A, and 1A. We investigated the 2 kb fragments upstream of all three TSSs. We refer to these fragments as *cis*-regulatory elements (CREs) A, B, and C, which together contain 42 fixed differences between ZAL2 and ZAL2^m^ (0.7% divergence; Fig. 4A; Table S7)^20^.

**Figure 4.**
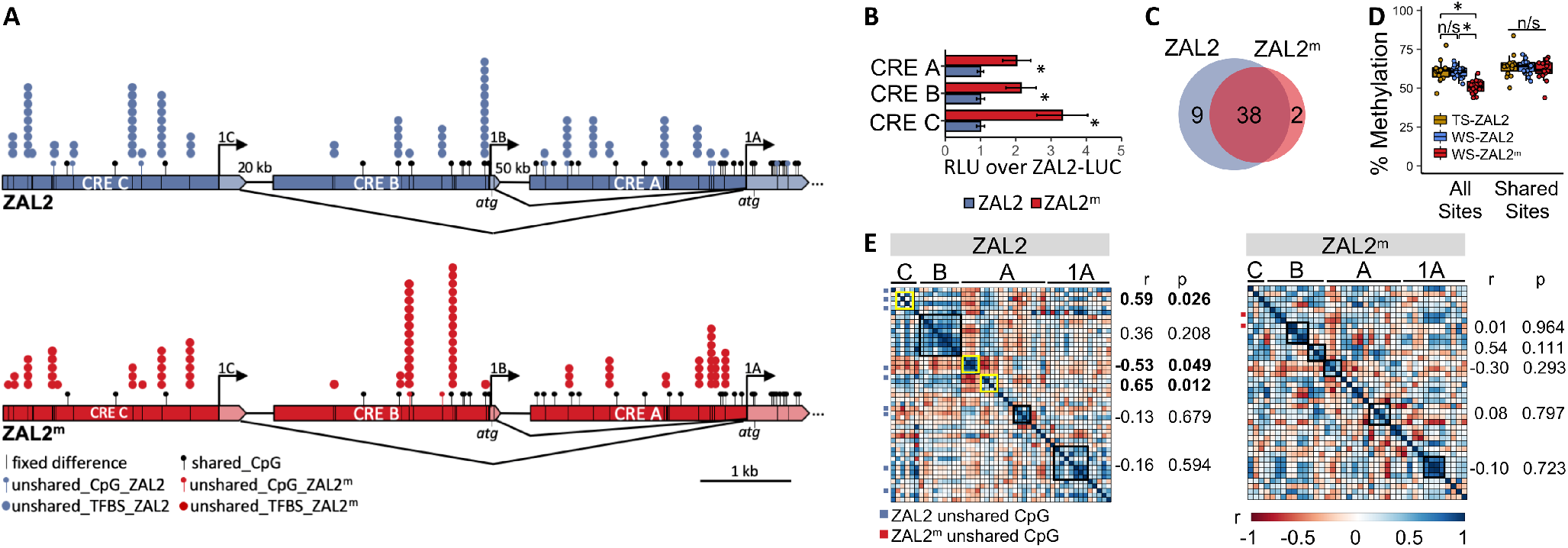
Mechanisms that potentially contribute to AI in *ESR1*. (A) *ESR1* is alternatively spliced. Dark blue or red regions are *cis*-regulatory elements (CREs); transcribed regions are light colors. The black lines represent 42 fixed differences that distinguish the ZAL2 from the ZAL2^m^ allele. Lollipops represent CpG sites. Stacked circles above the gene diagrams represent the transcription factors that are expressed in TnA and for which a binding site is disrupted by a fixed difference. (B) Bar graphs show the contribution of *cis*-regulatory variation in *ESR1* to variation in cis-regulatory element (CRE) activity in avian cells *in vitro*. The activity, in relative light units (RLU), of the ZAL2^m^-luciferase (LUC) (red) and ZAL2-LUC (blue) constructs are shown normalized to activity of the ZAL2-LUC construct. * p < 0.05. (C) Together, the *ESR1* CREs and exon 1 contain 49 CpGs: 38 are shared between the alleles, two are ZAL2^m^-specific and nine ZAL2-specific. (D) In TnA, methylation of these CREs depends on allele (p < 0.0001). In WS birds, the ZAL2^m^ sequence is less methylated than ZAL2 (p < 0.001). The ZAL2 sequences are not differentially methylated in WS compared with TS birds (p = 0.933; Table S10). Differential methylation of ZAL2^m^ and ZAL2 is not detectable when considering only shared CpG sites (p = 0.831). (E) Correlation matrices, using data from WS birds, demonstrate covariation in methylation of CpG sites along the sequences. Similarly methylated clusters that significantly predicted allele-specific expression are outlined in yellow; those that did not are outlined in black (see Fig. S6). Unshared sites are marked by blue (ZAL2) or red (ZAL2^m^) boxes to the left of each matrix to indicate which allele contains the site. * p < 0.05.

To test whether these differences are sufficient to cause allelic differences in transcription activity, we performed luciferase reporter assays using the ZAL2 and ZAL2^m^ alleles of each CRE (Fig. 4B, S4). We performed these assays in DT40 cells, which are one of the only available avian cell lines amenable to transfection.^21,22^ For all three CREs, the activity of the ZAL2^m^ allele was higher than the ZAL2 allele (Fig. 4B; Tables S8, S9). This result suggests that the genetic differentiation between the alleles is large enough to cause differential gene expression even in the absence of other factors (e.g., differential *trans-regulatory* elements or chromatin status) that may contribute to morph differences in expression.

To explore the impact of allelic differentiation on transcription factor binding, we identified nearly 300 transcription factor binding sites on ZAL2/ZAL2^m^ that are disrupted by SNPs^20^ – meaning that a SNP either abolished the site or reduced its affinity for a particular transcription factor, compared with the other allele. After identifying the transcription factors associated with each of these allele-specific binding sites, we used RNA-seq data to determine the availability of each factor in TnA. This approach revealed 120 transcription factors that are both associated with allele-specific binding sites and are expressed in TnA in both morphs (Fig. S5), suggesting a clear mechanism by which the level of *ESR1* expression could be strongly influenced by genotype. These transcription factors were neither overrepresented on ZAL2^m^ nor enriched for differential expression by morph (Fig. S5).

Differential gene expression and AI do not always involve genetic differentiation. These phenomena can also be caused by epigenetic factors, such as DNA methylation. Of 49 CpG sites in the regulatory sequences of *ESR1* (Fig. 4A), 22% are “unshared”, meaning that they contain a SNP and thus are present on only one of the two alleles. The number of polymorphic CpGs is higher on ZAL2 (Fig. 4C; Table S10), thus genetic divergence has affected CpGs, the primary targets of DNA methylation. In the next series of studies, we tested whether the regulatory regions of the two *ESR1* alleles are differentially methylated and whether that methylation

 predicts expression. We sequenced bisulfite-converted DNA extracted from TnA and compared the average percent methylation of the three *ESR1* CREs and exon 1A between the ZAL2 and ZAL2^m^ alleles. We found that the sequences were significantly more methylated on the ZAL2 than on the ZAL2^m^ allele (p < 0.001; Fig. 4D; Table S11), suggesting the morph difference in expression could be due in part to epigenetic control. We next tested the extent to which this differential methylation could be attributed to differentiation of genetic sequences. Indeed, when we considered only shared sites, in other words CpG sites present on both alleles, we could no longer detect differential methylation (Fig. 4D). Methylation of shared sites was largely the same between ZAL2 and ZAL2^m^ in WS birds. Further, methylation of the ZAL2 allele was essentially equivalent in the WS and TS birds (Fig. 4D). Therefore, any differential methylation that contributes to morph differences in expression is likely occurring mostly at genetically differentiated CpG sites, not shared sites.

We next wanted to test whether methylation of the CREs, particularly at unshared sites, predicts expression for each allele. As a means of data reduction, we constructed correlation matrices of CpG methylation to identify “clusters” of similarly methylated, neighboring sites within each allele (Fig. 4E). These clusters varied according to allele (Fig. 4E), suggesting that genetic differentiation has likely both contributed to and disrupted interactions between neighboring sites. The clusters that predicted allele-specific expression were found in CRE C and the 5’ end of CRE A, distal from exon 1A (Fig. S6). Remarkably, of the three clusters that significantly predicted allele-specific expression, all contained at least one unshared CpG site. Further, all of the predictive clusters were located on ZAL2, not on ZAL2^m^. In other words, the predictive value of methylated CpG sites was markedly reduced for ZAL2^m^, which is missing the ZAL2-specific sites. Overall, our findings suggest that methylation of ZAL2-specific sites is an important regulator of *ESR1* expression. Notably, this epigenetic regulation can be attributed predominantly to genetic differentiation between the alleles. Thus, this system represents a rare and interesting example of *cis*-regulation that is attributable to a combination of genetic and epigenetic forces.^23,24^ The regulatory variation in *ESR1* predicts and likely causes differential expression of this gene, which as we showed in the previous sections, can drive divergent behavioral phenotypes.

## Discussion

Here, we have demonstrated through a series of studies how genetic divergence in a single gene has contributed to the evolution of an aggressive phenotype in a naturally occurring, wild species of vertebrate. The gene *ESR1* has been captured by a chromosomal rearrangement that comprises a supergene, in other words a group of linked genes that are co-inherited. Supergenes caused by inversions are associated with social behavior not only in white-throated sparrows but also in ruffs, Alpine silver ants, and fire ants.^25–27^ The genes captured inside these inversions are in tight linkage disequilibrium, making it difficult to identify causal alleles or to uncover the genetic and epigenetic mechanisms that affect their expression. Here we show that identifying such alleles is possible when genomic resources are available, the model is amenable to mechanistic, experimental approaches, and there is already some existing knowledge about the physiological mechanisms underlying the behavior—in this case, steroid hormones.

Our results contribute to a growing literature indicating that hormone receptors and their ligands may be especially poised to play an important role in behavioral evolution.^28–31^ Given the role of ERα as a transcription factor, this gene is well-positioned to regulate gene networks thought to contribute to territorial aggression.^17,32^ The rapid effects of E2 on aggression that we detected (Fig. 2C,F) are likely attributable to ERα situated in cell membranes.^33^ In rats, membrane-bound ERα interacts with a variety of other molecules, for example a metabotropic glutamate receptor mGluR;^34^ in white-throated sparrows, the gene encoding this glutamate receptor is also inside the ZAL2^m^ supergene, as are genes encoding a number of other factors that interact with ERα.^6,17^ Thus, *ESR1* is likely to be co-evolving with other genes inside the supergene. The ZAL2/2^m^ genomic architecture favors the inheritance of co-adapted alleles, which is thought to be important for the evolution of complex phenotypes.^35^ Overall, this species offers promising new avenues for understanding co-adapted gene complexes and their role in behavior.

## Methods

### Effects of ESR1 knockdown on aggression

#### Non-breeding white-throated sparrows

We collected 31 white-throated sparrows in mist nets on the campus of Emory University in Atlanta, GA during fall migration, when gonads are regressed and plasma levels of sex steroids are low^36,37^. Our rationale for testing birds in non-breeding condition was that (1) the morph difference in *ESR1* expression in TnA is pronounced in non-breeding birds^13^ and (2) ERα would be relatively unoccupied and thus manipulatable with exogenous E2^11^. Sex and morph were confirmed by PCR^38,39^. The PCR to determine morph amplifies an indel on chromosome 2. We follow conventional nomenclature for avian chromosomes, numbering them from largest to smallest^40^; chromosome 2 in white-throated sparrows corresponds to chromosome 3 in chicken^39^. Birds were housed in the Emory animal care facility in walk-in flight cages (4’ x 7’ x 6’), supplied with *ad libitum* seed and water. The day length was kept constant at 10L:14D, to prevent spontaneous gonadal recrudescence. At least one month prior to behavioral assays, birds were transferred to individual cages (15” x 15” x 17”) inside identical walk-in sound-attenuating booths (Industrial Acoustics, Bronx, NY), two to six birds per booth. In order to habituate birds to the presentation of wax moth larvae (*Achroia grisella*) as food items, which would be used in the E2 manipulation, each bird received one larvae per day. Birds that consistently and reliably ate the larva within one min of presentation were included in the study.

#### Cannulation surgeries

Each bird was fitted with bilateral 26-gauge, stainless steel guide cannulae aimed at TnA. Surgeries were performed using a stereotaxic apparatus with ear bars and beak holder custom-designed for birds. To place the cannulae we used a custom-designed holder that accommodates two cannulae with a fixed distance of 3.8 mm between them (PlasticsOne, Roanoke, VA). Four animals were used to establish stereotaxic coordinates and proper angles of approach for TnA. Final coordinates, relative to the anterior pole of the cerebellum, were AP 0.0 mm, ML 1.9 mm. Cannulae were lowered to a depth of 3.6mm and mounted to the skull using dental acrylic and veterinary-grade tissue adhesive (VetBond; 3M). A 33-gauge stainless steel obturator, which extended 1 mm beyond the tip of the guide cannula, was inserted on each side. Birds recovered from surgery for at least 3 days before dominance testing (see next section).

#### Pre-screening for social dominance

To quantify aggression, we presented each focal animal with a subordinate “opponent” in an adjacent cage. In order to ensure that each opponent was subordinate to the focal animal, we pre-screened the birds in dyads to determine their dominance relationships. This screening was performed according to Merritt et al.^11^. Briefly, during prescreening trials, we placed the cages of two same-morph, cannulated birds adjacent to one another in an otherwise empty sound-attenuating booth equipped with a video camera ~1m from their cages. We recorded their interactions for 30 min, then returned each bird to its home booth. Dominance was operationalized according to Merritt et al.^11^ An observer scored the videos for two behaviors: attempted attacks by both birds, defined as the bird making contact with both feet on the wall of its cage facing the other bird and flapping its wings, and the amount of time (s) each bird spent in proximity to the other cage. Proximity was defined as being within an area ~12cm from the wall of the cage closest to the other bird (Fig. 2B). This area was marked on each focal animal’s cage and visible in the videos. The member of the dyad that made more attacks and spent more time (s) in proximity to the other bird’s cage was deemed dominant. In the behavioral trials described below, the dominant bird was used as the focal animal and the subordinate was the opponent. If neither bird in the dyad dominated in terms of both behaviors, the dyad was dissolved and each bird was tested again with a new bird. Most dyads were same-sex; however, some dyads consisted of a male focal animal and a female opponent. No female engaged in copulation solicitation or other courtship behavior during testing, and whether the dyad was same-sex or mixed-sex did not affect the behavior of either the focal animal (attacks F = 0, p = 0.985; time F = 0.51, p = 0.484) or the opponent (attacks F = 0.08, p = 0.78; time F = 1.68, p = 0.21). We emphasize that all birds were in non-breeding condition and do not engage in courtship behavior during the nonbreeding season.

#### Antisense

*ESR1* expression was manipulated with custom-designed locked nucleic acid (LNA^™^) 15-mer antisense oligonucleotides designed by Qiagen following Kelly & Goodson.^41^ The antisense oligo for *ESR1* knockdown (*ESR1*-KD; TTCAAAGGTGGCACT) was designed to target the stop codon of the *ESR1* transcript (XM_026794125.1), avoiding ZAL2/ZAL2^m^ SNPs, starting 11 bp upstream of the stop codon. A scrambled control oligonucleotide was designed from the same nucleotides, but in a random order (TAGCATGTCAGATCG). Both the *ESR1*-KD and scrambled sequences were searched on BLAST (NCBI) against the *Zonotrichia albicollis* refseq_rna to confirm no significant alignments to other transcripts, in particular *ESR2* (XM_014270428.2).

Starting the day after the establishment of a dyad, we made a series of four injections of antisense or scrambled oligos (1 μg in 0.25 μl isotonic saline), 2 per day, 10 hrs apart (within the first and last hour of light of the light/dark cycle). 250 nl microinjections were made slowly (2 min per side to avoid tissue damage) using a Hamilton neurosyringe connected to a 33 gauge injector via ~20 cm of polyethylene tubing. Cannulae were checked for leakage after each infusion and none was noted.

#### Hormone manipulation

We administered exogenous E2 via a non-invasive, minimally stressful method that increases plasma E2 to a similar, high level in both morphs^11^. Before behavioral testing (below), a larva of the wax moth was injected with either 300 μg of cyclodextrin-encapsulated 17β-estradiol (E2; Sigma-Aldrich, cat. no. E4389) or cyclodextrin alone as a control (CON; Sigma-Aldrich, cat no. C0926) using a Hamilton syringe. The injected larva was then fed to the focal animal. This oral dosage of E2 increases plasma E2 to a similar extent in WS and TS birds, and produces levels typical of breeding WS females.^11^

#### Behavioral testing

Behavioral testing was conducted the day after the fourth injection, as previously described^11^. Briefly, before a behavioral trial, we placed the focal bird in its cage in an empty sound-attenuating booth to acclimate it to that environment. After one hr, we placed the opponent, in its cage, immediately adjacent to the focal bird’s cage. An opaque barrier visually isolated the birds from each other. At the same time, a wax moth larva that had been injected immediately prior with E2 or CON was placed on the floor of the focal bird’s cage and the experimenter immediately left the room. The birds were then monitored remotely to determine exactly when they consumed the larvae.

Ten min after the focal bird consumed the larva, the opaque barrier was removed and the birds were allowed to interact for 10 min. Attacks and time spent in proximity to the opponent were scored in de-identified videos, as described above for the pre-screening dominance trials, by an observer blind to morph (which is not easily assessed in videos), oligo type (antisense or scrambled), and hormone treatment (E2 or CON). As was also reported by Merritt et al. and Heimovics et al.,^11,33,42^ singing occurred too infrequently for statistical analysis. After a washout period of 48 hrs, each focal bird participated in a second trial with the opposite hormone treatment (CON or E2). All trials were counterbalanced with respect to the order of treatment.

Data were analyzed as described by Merritt et al.^11^ using generalized linear models (GLM) with a Gaussian distribution. The best-fit model was selected on the basis of AIC values.^43^ Wald chi-squared tests were used to generate analysis-of-deviance summary tables. These analyses were performed with the function *glm* from the *lme4* (v. 1.1-15) package^44^ and summarized using the function *Anova* from the *car* (v.3.0-3) package.^45^ All analyses included the fixed effect of minute (1-10, over the course of the 10-min trial) and the random effect of individual.

#### Verification of cannula placement

Birds were euthanized by isoflurane overdose (Abbott Laboratories, North Chicago, IL) 24 hrs after the second behavioral trial. Each cannula was then injected with bromophenol blue. Brains were dissected from the skulls, frozen rapidly on dry ice, and sectioned on a cryostat in order to verify cannula placement and dissect tissue for qPCR. We then punched and extracted RNA (see Supplemental Methods), such that punches were made at the site of dye, one per hemisphere, for a total of 2 punches per bird. We considered birds with no dye in TnA to be misses (n = 7), and in these cases an additional 2 punches were made in TnA. Misses were determined by an observer blind to morph and oligo type (antisense or scrambled). The final sample size for each group, each of which included multiple males and females, was WS *ESR1*-KD n = 6, WS scrambled n = 7, TS *ESR1*-KD n = 6, TS scrambled n = 7.

cDNA was produced by reverse transcription using the Transcriptor First Strand cDNA synthesis kit with random hexamer primers, then the reaction was diluted to 20 ng/ul for qPCR. We designed exon-spanning primers to be used with probes from the Roche Universal Probe Library ^46^ for *ESR1*^12^ (F: GCACCTAACCTGTTACTGGACA; R: TGAAGGTTCATCATGCGAAA; Probe 132) and *ESR2* (F: GAAGCTGCAGCACAAGGAGT; R: CCTCTGCTGACCAGTGGAAC; Probe 151). The amplified sequence for *ESR2* was verified via agarose gel and Sanger sequencing. qPCR was performed using a Roche LightCycler 480 Real-Time PCR System in triplicate for each sample on 384-well plates as previously described^47^. Using the LightCycler 480 Software Version 1.5.0, we calculated crossing point (Cp) values using the Abs Quant/2nd Derivative Max method. We normalized the expression of each gene of interest to two reference genes: peptidylprolyl isomerase A (*PPIA*) and glyceraldehyde-3-phosphate dehydrogenase (*GAPDH*), which have been previously validated for use in white-throated sparrow brain tissue^47^. We performed GLMs, as chosen on the basis of AIC values,^43^ to test for effects of morph and oligo type (antisense or scrambled) on the expression of *ESR1*, and then followed up the significant interaction with GLMs testing for an effect of oligo type on expression of *ESR1* within each morph. Wald chi-squared tests were used to generate analysis-of-deviance summary tables. We used Pearson’s partial correlations to test whether the degree to which E2 facilitated aggression (E2-CON) could be explained by the level of *ESR1* or *ESR2* expression, controlling for morph, using function *pcor.test* from the package *ppcor*^48^ (v. 1.1). We also asked whether *ESR1* and *ESR2* expression were correlated with each other using a Pearson’s correlation *cor.test* in base R.

### Quantification of *ESR1* allelic imbalance

#### Collection of free-living white-throated sparrows

Adults and nestlings of both sexes and morphs were collected at Penobscot Experimental Forest near Orono, ME. Adults were collected during the peak of territorial behavior, after pair formation and territory establishment but prior to or early in the stage of incubation.^3,17^ Before collecting the adults, we characterized their behavioral responses to territorial intrusion by conducting STIs (see Supplemental Materials).^49^ Birds remained on their territories for at least 24 hrs before we returned to collect tissue, in order to minimize the effects of the STI itself on gene expression. Later, during the parental phase of the breeding season, we collected nestlings on post-hatch day seven, two days before natural fledging.^16,50^ See Supplemental Material for details on tissue collection.

#### Microdissection

Our methods for quantifying *ESR1* expression in micropunched tissue have been described elsewhere.^12^ Briefly, we cryosectioned brains at 300 μm in the coronal plane, then used the Palkovits punch technique to obtain 1 mm diameter samples from the regions of interest as described by Grogan et al. and Zinzow-Kramer et al.^12,17^ TnA was sampled in each hemisphere in two consecutive sections for a total of four punches, which were pooled for nucleic acid extraction. For HYP, punches were centered on the midline such that they contained tissue from both hemispheres. One punch was made immediately ventral to the anterior commissure and included the caudal portion of the medial preoptic area, the paraventricular nucleus, and the anterior hypothalamus. A second punch was made ventral to the first and included the ventromedial hypothalamus. Both HYP punches were made in two consecutive sections for a total of four punches, which were pooled for nucleic acid extraction in the adult samples. Punches of the ventral and dorsal hypothalamus were kept separate for chicks and expression was averaged during data analysis. For rostral POM, one punch was made underneath the septo-mesencephalic tract and above the supraoptic decussation. See Supplemental Methods for RNA/DNA extraction.

#### Allelic imbalance assay

To detect AI, primers and allele-specific probes for the ZAL2 and ZAL2^m^ alleles of *ESR1* were designed by Integrated DNA Technologies (Coralville, Iowa, USA) to target a SNP within *ESR1* exon 1A (see Supplemental Materials). Only WS birds were used because TS birds do not have the ZAL2^m^ allele. Our sample size was as follows: for nestlings, HYP n = 26, TnA n = 27, POM n = 27; for adults, HYP n = 18, TnA n = 18, POM n = 15. Each group comprised roughly equal numbers of males and females. We then calculated relative expression within each reaction, averaged across three replicate reactions, and calculated the ZAL2^m^/ZAL2 expression ratio. The ratio for each cDNA sample was then normalized to the average of the ratios calculated from WS genomic DNA (gDNA) control samples (mean = 0.99, min = 0.95, max = 1.02) to correct for the relative affinity of each probe to its target sequence. We tested for AI within each region and age using one-sample t-tests comparing to a null ratio of 1. To determine whether the degree of AI varied by age or region, we analyzed the data in a two-way mixed ANOVA with region and age as factors (Table S5), followed by Tukey’s Honest Significant Difference (Table S6). Associations between AI and territorial responses were tested using Pearson’s correlations.

### Analysis of cis-regulatory variation in *ESR1*

#### Analysis of transcription factor binding sites

We examined the CREs in *ESR1* to identify transcription factor binding sites that are disrupted by ZAL2/2^m^. To predict differential transcription factor binding between the two alleles, we used the transcription factor affinity predictor tool for SNPs (sTRAP;^24,51^ See Supplemental Methods). We then entered the list of predicted differential sites into TRANSFAC version 2019.2 to create a comprehensive list of associated transcription factors predicted to bind with greater affinity to one allele than the other. We then used our RNA-seq data from TnA to identify the transcription factors that are expressed.^17^ Reads were counted and normalized in DESeq2 (v1.24.0)^52^. Finally, we cross-referenced the list of factors predicted to bind differentially to the two alleles with the list of factors that are expressed in TnA.

#### Luciferase assays

A 2 kb sequence upstream of each TSS of *ESR1* was amplified by PCR using gDNA from a WS (ZAL2/ZAL2^m^) bird, then cloned upstream of firefly luciferase into the pGL3-control vector at the KpnI and MluI restriction sites (Fig. S4C; Table S12). Clones containing the ZAL2 or ZAL2^m^ alleles were identified, on the basis of known fixed differences, via Sanger sequencing by Genscript (Piscataway, NJ, USA). Luciferase reporter assays (see Supplemental Materials) were performed in three different cell types: HeLa, HEK-293, and DT40 cells. Twenty-four hrs after transfection, luciferase activity was quantified using the Dual-Glo assay system (Promega) on a Biotek Synergy plate reader. The value for the ZAL2^m^ allele was normalized to the value for the ZAL2 allele, meaning that the ZAL2^m^ data were expressed relative to a value of 1 for the ZAL2 allele (Fig. 4B). Effects of CRE, allele, and interactions between CRE and allele were assessed using a 2-way ANOVA followed by Student’s pairwise t-tests.

#### Analysis of DNA methylation

We bisulfite-converted 200 ng of gDNA, extracted from the punches from TnA of the adults described above (see *Microdissection*), using the EZ DNA Methylation-Lightning Kit following the manufacturer’s instructions (Zymo, Irvine, CA, USA). Thirteen amplicons containing shared and unshared CpG sites in the three CREs and exon 1A of *ESR1* were amplified using PCR with primers that do not fall on SNPs or CpGs (Table S12; See Supplemental Methods). All 13 amplicons for each sample were pooled and 5 μl of that pool was used for next-generation sequencing. Adapter-ligated libraries were constructed using the 16S Metagenomic Sequencing Library kit (Illumina; San Diego, USA) following the manufacturer’s instructions. Samples were then run on the Agilent bioanalyzer to confirm successful indexing and quality of total DNA, then pooled for a single sequencing run (PE300) on the Illumina MiSeq at the Emory Integrated Genomics Core.

Reads were filtered, trimmed and aligned to a bisulfite-converted, N-masked reference genome (see Supplemental Materials). Reads were then filtered for non-CpG methylation such that any read that included 3 C’s in a row in non-CG positions was removed. Reads were then assigned to alleles on the basis of non-CpG fixed SNPs using SNPsplit (v0.3.4).^53^ Bismark^54^ then extracted methylation calls and generated bedGraphs that were imported into RStudio for further analysis. CpGs were filtered for low coverage (<10x) and analyzed using a custom script in R. To arrive at an overall level of methylation for each of the two alleles for each bird, we averaged % methylation across all of the CpGs in that allele. When including unshared CpG sites, we treated that site on the other allele as 0% methylation. We then ran linear regressions with allele as a single factor with three levels, TS-ZAL2, WS-ZAL2, and WS-ZAL2^m^, and with bird as a random factor, using the *lme4* (v1.1-21) package^44^. These results were summarized using the *car* (v3.0-3) package^45^ and post-hoc tests (estimated marginal means controlling for individual) were conducted using the *emmeans* (v1.4.3.01)^55^ package. We then created Pearson correlation matrices to identify clusters of significantly correlated CpGs within each allele. We averaged % methylation across the CpGs within each cluster (six clusters on ZAL2, five on ZAL2^m^) following established protocols.^56,57^ We then used Pearson correlations to calculate the extent to which the % methylation of each cluster predicted allele-specific expression in our RNA-seq data. For males (n = 8 TS, 10 WS), we used normalized read counts from previously published data (11,12). For females (n = 6 TS, 6 WS), we used normalized read counts from new data (See Supplemental Materials for sequencing methods).

## Author Contributions and Notes

J.R.M, K.E.G, W.M.Z-M, E.A.O, S.V.Y, and D.L.M designed research, J.R.M, K.E.G, W.M.Z-M, D.S., E.A.O, S.V.Y, and D.L.M performed research, J.R.M, K.E.G, W.M.Z-K, D.S, E.A.O, S.V.Y, and D.L.M analyzed data, and J.R.M and D.L.M wrote the paper.

The authors declare no conflict of interest.

This article contains supporting information online.

## Acknowledgments

We are grateful to Carlos Rodriguez-Saltos, T.J. Libecap, Suzanne Mays, William Hudson, Emily Kim, and Aubrey Kelly for technical assistance. This work was supported by 1R01MH082833 to D.L.M, NSF 16-505 to D.L.M, and 1F31MH114509 to J.R.M.

