## Supplemental materials for "A behavioral polymorphism caused by a single gene inside a supergene" (from another version of this preprint: version 1, 2020-01-14)

### Supplemental methods

All procedures involving animals were approved by the Emory University Institutional Animal Care and Use Committee, were in keeping with all federal, state, and local laws, and adhered to guidelines set forth by the National Institutes of Health Guide for the Care and Use of Laboratory Animals. All data were analyzed in RStudio (v1.2.1335; R v3.6.0). The $\alpha$ level was set at $p \leq 0.05$ unless otherwise specified.

*Simulated territorial intrusions (STI) and collection of tissue from breeding white-throated sparrows.* STIs were performed on free-living, territorial white-throated sparrows on two consecutive days at the same time each day. We presented either a TS or WS live male decoy in a counterbalanced order.[1] During presentation of the decoy, conspecific song was played for ten min. Aggressive responses, including songs, were quantified for both the male and the female resident. At least 24 hr after the second STI, we returned to the site to collect both the male and the female. We used song playback to lure them quickly into a mist net. Average latency to capture was 6.1 ± 0.89 SE min after the onset of the song playback. There were no effects of sex ($F = 0.167$, $p = 0.602$) or morph ($F = 0.678$, $p = 0.686$) on time to capture, and no interaction between sex and morph ($F = 0.138$, $p = 0.713$). All birds were sacrificed immediately after capture by isoflurane overdose. Brains were rapidly dissected from the skulls, frozen on dry ice and stored at -80 °C. Sex and morph for all birds were later confirmed by PCR.[2,3]

*RNA/DNA extraction.* DNA and RNA were extracted using either the Qiagen Allprep RNA/DNA micro kit (Qiagen, Valencia, CA, USA) with modifications or the Qiagen miRNeasy micro kit (Qiagen, Valencia, CA, USA) with modifications (see Zinzow-Kramer et al.[4,5]). The DNA was used for the bisulfite conversion, see below. The RNA was reverse transcribed into cDNA using the Transcriptor First Strand cDNA synthesis kit with random hexamer primers (Roche Diagnostics, Indianapolis, IN, USA). cDNA reactions were then diluted to 20 ng/ul for qPCR.

*Allelic imbalance assay.* To detect allelic imbalance (AI), primers (F: GCAGGATTTCACTCCCTGAA, R: TACCTGTTGGCTGTGATGATG) and allele-specific probes for the ZAL2 (CTGTC**G**CCCT) and ZAL2$^m$ (CTGTC**A**CCCT) alleles of *ESR1* were designed by Integrated DNA Technologies (Coralville, Iowa, USA) to target a SNP (NW_00508196.1:1801511) within ESR1 exon 1A following Chen et al.[6] To increase the melting temperatures and thus specificity, the probes contained 4-6 locked nucleic acids. The probes were labeled on the 5' end with dye; 6-FAM was used for ZAL2 and 5Cy5 for ZAL2$^m$. All samples were run in triplicate in on a Roche LightCycler 480 Real-Time PCR System. In addition to experimental samples, each plate included the following controls: a negative cDNA reaction control with no template added, a negative cDNA reaction control with no RT enzyme added, positive controls of ZAL2/ZAL2 and ZAL2$^m$/ZAL2$^m$ genomic DNA, and a positive control with a 5x dilution of WS gDNA to assess overall amplification efficiency. We also included a dilution curve from 1:8 to 8:1 of ZAL2: ZAL2$^m$ genomic DNA to assess the specificity of the probe binding (Fig. S2B). To correct for imperfect specificity of the probes, we also included >10 undiluted ZAL2/ZAL2$^m$ genomic DNA samples on each plate. Each reaction contained approximately 5-25 ng of cDNA depending on the brain region, 750 nM of forward and reverse primer, 6.0 ul of 2X Probes Master (Roche), and 200 nM of each probe. Cycling conditions were 95° C for 10 min, followed by 45 cycles of 95° C for 10 seconds, 60° C for 30 seconds, and 72° C for 1 second. We removed a maximum of one technical outlier from each triplicate (st. dev. > 0.2). Samples that produced hyper-variable results or for which the reaction failed were not included in final analyses (final sample size for chicks: HYP n = 26, TnA n = 27, POM n = 27; adults: HYP n = 18, TnA n = 18, POM n = 15). We then used the Abs Quant/Fit Points method in the LightCycler 480 Software Version 1.5.0 to calculate relative expression from the crossing point (Cp) values, averaged across the replicate reactions, then divided ZAL2/ZAL2$^m$. The primer efficiency was ~1.8 and the ratio of triplicate averages of each allele (cDNA) was normalized to the average of the ratios calculated from all of the gDNA samples (mean = 0.99, min = 0.95, max = 1.02) to correct for incomplete probe binding.

*sTRAP*. We submitted 42 ZAL2 and ZAL2$^m$ sequences of 120 bp each, centered at each fixed sequence difference (41 SNPs and one indel), to the transcription factor affinity predictor tool for SNPs sTRAP.[7] We selected “transfact.12.1 metazoans” with the “chordate_conserved_elements” background model. sTRAP computed the affinity of transcription factors for each allele, then ranked the difference in p-values between the alleles ($\log_{10}$(ZAL2 p-value/ZAL2$^m$ p-value)). We corrected for multiple comparisons using Benjamini-Hochberg corrections. Position weight matrices with significant ($p < 0.05$) affinity for at least one allele and a p-value ratio >1.5 were considered by sTRAP to be differential binding sites.

*Luciferase assays*. Two hundred ng of each construct was co-transfected into cells, along with 10 ng of constitutively expressed Renilla luciferase, using FuGene HD (Promega) in OptiMEM. HEK-293 cells were cultured in DMEM supplemented with 10% fetal bovine serum (FBS, Atlanta Biologicals). HeLa cells were cultured in MEM$\alpha$ supplemented with 10% FBS. DT40 cells were cultured in DMEM with 10% FBS and 5% chicken serum and supplemented daily with 0.05 mM 2-mercaptoethanol following ATCC guidelines (https://www.atcc.org/products/all/CRL-2111.aspx). Data collection was completed separately for each cell type as follows: 24 hr after transfection, the activities of firefly and Renilla luciferase were read three times using the Dual-Glo assay system (Promega) on a Biotek Synergy plate reader. These three readings were then averaged for firefly luciferase as well as for Renilla luciferase. The average reading for firefly luciferase was normalized to that of Renilla luciferase for each well. This normalized value was then averaged across five replicates. The resulting value for the ZAL2$^m$ allele was then normalized to the value for the ZAL2 allele, meaning that the ZAL2$^m$ data were expressed relative to a value of 1 for the ZAL2 allele (Fig. 4B). Experiments were replicated three times for the HeLa and HEK cells and six times for the DT40 cells; results were averaged across experiments. Effects of CRE (A, B, or C), allele, and the interaction between CRE and allele were assessed using a 2-way ANOVA. When significant effects or interactions were found, we used Student’s pairwise t-tests to test for the effect of allele within each CRE.

*PCR of bisulfite-converted DNA*. Each 25 µl PCR included 0.5 µl JumpStart Taq DNA polymerase, 2.5 µl JumpStart buffer containing 40 mM $MgCl_2$, 5 nM of each primer (Table S12), and 8 ng of bisulfite-converted DNA. The PCR protocol for each primer pair was optimized using a temperature and $MgCl_2$ gradient such that an additional 0-100 mM $MgCl_2$ was added to some reactions. The PCR protocol was: (1) 60 s at 95°C, (2) 30 s at 95°C, (3) 30 s at 55°C, (4) 60 s at 70°C, and then (5) 5 min at 70°C for a final extension phase, repeating steps 2-4 for 40 cycles. Ten µl of each PCR product was visualized on a 1% agarose gel. Any PCRs that did not show a band were rerun, modifying the concentration of $MgCl_2$, until a band of the correct size was seen.

*bsDNA filtering and mapping.* We filtered reads at a Phred score of 30, then trimmed for low quality, Nextera adapter sequences, and indexes using Trim-galore! (v0.3.7).[8] A bisulfite-converted reference genome was generated in Bismark (v0.18.1)[9] using the scaffold that contains *ESR1* (NW_005081596.1) from the genome of a TS bird (GCF_000385455.1). This reference genome was N-masked at ZAL2/ZAL2$^m$ fixed SNPs[10] in order to reduce mapping bias, and unshared ZAL2$^m$ CpGs were added to the reference genome so that information on methylation state could be extracted for the highest possible number of CpGs. Bismark does not use CpGs to map reads, so this modification did not introduce bias into our analysis. Reads were aligned using Bismark with bowtie2 (v2.3.5).[11]

*RNA-seq library preparation, sequencing, and data analysis.* Library preparation and sequencing of six TS females and six WS females were performed by the Duke Center for Genomic and Computational Biology Sequencing and Genomic Technologies Core in Durham, NC. After assessing the RNA quality using Qubit and Agilent Tapestation (Agilent Technologies, Santa Clara, CA, USA), RNA-seq libraries were constructed using a Kapa Stranded mRNA-Seq Kit (Kapa Biosystems, Boston, MA, USA) following the manufacturer's instructions. The samples were then pooled and sequenced on an Illumina HiSeq 4000 in pools of 8-10 samples per lane in 150 paired-end reactions. We obtained 20-45 million read pairs per sample. RNA-seq files were trimmed using Trim-galore! (v0.4.5)[8] retaining only pairs of sequences for which all bases had an average Phred score of 30. Again to control for mapping bias towards the reference (ZAL2/ZAL2) genome, we mapped all reads to the above-mentioned N-masked genome (for further details, see Sun et al.[10]). Reads were mapped with STAR (v2.5.2b)[12] using the 2-pass mode, and only uniquely mapped reads were retained for differential expression analysis. To obtain read counts of *ESR1* at the allele level, reads were assigned to ZAL2 or ZAL2$^m$ using SNPsplit (v0.3.3),[13] counted by htseq-count (v0.11.1),[14] and then normalized and analyzed using DeSeq2 (v1.24.0).[11]

**Table S1.** F and *p*-values for ANOVA accompanying Fig.2A in the main text.

| Omnibus ANOVA | *F* | *p* |
|---|---|---|
| Antisense | 4.46 | 0.049 |
| Morph | 8.82 | 0.008 |
| Antisense x Morph | 4.84 | 0.041 |

**Table S2.** *P*-values for post-hoc comparisons (Tukey's Honest Significant Difference) accompanying Fig.2A in the main text.

| | Post-hoc pairwise comparisons (*p*) | | |
|---|---|---|---|
| | Effect of Antisense within Morph | Effect of Morph by Antisense Treatment | |
| Morph | *ESR1*-KD - SCR | Antisense | WS-TS |
| TS | 0.994 | *ESR1*-KD | 0.947 |
| WS | 0.048 | SCR | 0.011 |

**Table S3**. $\chi 2$ and *p*-values for the GLMs accompanying Fig.2C and 2F in the main text.

| | | Time Spent Near the Opponent (s) | | Attacks | |
|---|---|---|---|---|---|
| SCR birds | | $\chi^2$ | *p* | $\chi^2$ | *p* |
| | Morph | 0.34 | 0.559 | 0.02 | 0.886 |
| | Treatment | 0.07 | 0.790 | 0.37 | 0.542 |
| | Minute | 5.74 | 0.017 | 22.69 | < 0.001 |
| | Morph x Treatment | 14.72 | < 0.001 | 8.00 | 0.005 |
| *ESR1*-KD birds | | | | | |
| | Morph | 0.00 | 0.996 | 6.14 | 0.013 |
| | Treatment | 0.56 | 0.456 | 11.24 | < 0.001 |
| | Minute | 0.44 | 0.509 | 14.02 | < 0.001 |
| | Morph x Treatment | 0.14 | 0.712 | 0.49 | 0.483 |
| WS birds | | | | | |
| | Antisense | 24.41 | < 0.001 | 9.64 | 0.002 |
| | Treatment | 5.03 | 0.025 | 0.24 | 0.627 |
| | Minute | 13.32 | < 0.001 | 20.21 | < 0.001 |
| | Antisense x Treatment | 4.43 | 0.035 | 9.53 | 0.002 |
| TS birds | | | | | |
| | Antisense | 15.53 | < 0.001 | 0.04 | 0.837 |
| | Treatment | 8.04 | 0.005 | 9.77 | 0.002 |
| | Minute | 0.00 | 0.976 | 16.19 | < 0.001 |
| | Antisense x Treatment | 4.46 | 0.035 | 1.21 | 0.271 |

**Table S4.** $\chi^2$ and *p*-values for effect of E2 within each antisense/morph group accompanying Fig.2C and 2F in the main text.

| | | Time Spent Near the Opponent (s) | | Attacks | |
|---|---|---|---|---|---|
| WS *ESR1*-KD | | $\chi^2$ | *p* | $\chi^2$ | *p* |
| | Treatment | 0.07 | 0.796 | 3.24 | 0.072 |
| | Minute | 6.29 | 0.012 | 5.44 | 0.020 |
| WS SCR | | | | | |
| | Treatment | 6.58 | 0.010 | 6.96 | 0.008 |
| | Minute | 8.35 | 0.004 | 16.59 | < 0.001 |
| TS SCR | | | | | |
| | Treatment | 7.48 | 0.006 | 9.81 | 0.002 |
| | Minute | 0.00 | 0.977 | 16.24 | < 0.001 |
| TS *ESR1*-KD | | | | | |
| | Treatment | 0.57 | 0.451 | 8.20 | 0.004 |
| | Minute | 1.17 | 0.279 | 8.55 | 0.003 |

**Table S5.** F and *p*-values for the omnibus test (ANOVA) accompanying Fig.3 in the main text.

| Omnibus ANOVA | *F* | *p* |
|---|---|---|
| Region | 82.86 | < 0.001 |
| Age | 13.54 | < 0.001 |
| Region x Age | 12.9 | < 0.001 |

**Table S6.** *P*-values for post-hoc comparisons (Tukey's Honest Significant Difference) and t, d.f., and *p*-values for allelic imbalance (1-sample t-test) accompanying Fig.3 in [illegible]

Post-hoc pairwise comparisons (*p*)

| | Region effects within Adults | | | Region effects within Chicks | | | Age effects within Region |
|---|---|---|---|---|---|---|---|
| Region | HYP | POM | TnA | HYP | POM | TnA | |
| HYP | - | 0.999 | < 0.001 | - | 0.997 | 0.059 | 0.988 |
| POM | 0.999 | - | < 0.001 | 0.997 | - | 0.019 | 0.999 |
| TnA | < 0.001 | < 0.001 | - | 0.059 | 0.019 | - | < 0.001 |

Allelic imbalance ($ZAL2^{m}$/ZAL2)

| | Adult | | Chick | |
|---|---|---|---|---|
| Region | *t* (d.f.) | *p* | *t* (d.f.) | *p* |
| HYP | 10.903 (15) | < 0.001 | 4.778 (25) | < 0.001 |
| POM | 8.071 (13) | < 0.001 | 12.569 (26) | < 0.001 |
| TnA | 4.739 (15) | < 0.001 | 2.616 (25) | 0.015 |

**Table S7.** Fixed SNPs and indels in the *ESR1 cis*-regulatory elements.

| Accession | Position | ZAL2/ZAL2$^{m}$ | CRE | Dist. from TSS |
|---|---|---|---|---|
| NW_005081596.1 | 1725878 | G/C | C | -1952 |
| NW_005081596.1 | 1725935 | C/T | C | -1895 |
| NW_005081596.1 | 1726061 | G/C | C | -1769 |
| NW_005081596.1 | 1726101 | C/G | C | -1729 |
| NW_005081596.1 | 1726110 | C/T | C | -1720 |
| NW_005081596.1 | 1726277 | T/C | C | -1553 |
| NW_005081596.1 | 1726287 | A/G | C | -1543 |
| NW_005081596.1 | 1726299 | C/T | C | -1531 |
| NW_005081596.1 | 1726315 | G/C | C | -1515 |
| NW_005081596.1 | 1726484 | G/T | C | -1346 |
| NW_005081596.1 | 1727022 | T/A | C | -808 |
| NW_005081596.1 | 1727091 | C/G | C | -739 |
| NW_005081596.1 | 1727273 | T/C | C | -557 |
| NW_005081596.1 | 1727539 | C/T | C | -291 |
| NW_005081596.1 | 1747958 | T/A | B | -1456 |
| NW_005081596.1 | 1747993 | A/G | B | -1421 |
| NW_005081596.1 | 1748584 | T/C | B | -830 |
| NW_005081596.1 | 1748651 | T/G | B | -763 |
| NW_005081596.1 | 1748701 | G/A | B | -713 |
| NW_005081596.1 | 1748949 | A/G | B | -465 |
| NW_005081596.1 | 1749043 | A/G | B | -371 |
| NW_005081596.1 | 1749073 | TA/T | B | -341 |
| NW_005081596.1 | 1749079 | A/T | B | -335 |
| NW_005081596.1 | 1749359 | C/G | B | -55 |
| NW_005081596.1 | 1749401 | C/T | B | -13 |
| NW_005081596.1 | 1799245 | C/A | A | -1857 |
| NW_005081596.1 | 1799456 | C/A | A | -1646 |
| NW_005081596.1 | 1799677 | A/T | A | -1425 |
| NW_005081596.1 | 1799821 | T/A | A | -1281 |
| NW_005081596.1 | 1800307 | G/A | A | -795 |
| NW_005081596.1 | 1800338 | T/C | A | -764 |
| NW_005081596.1 | 1800343 | A/G | A | -759 |
| NW_005081596.1 | 1800344 | A/C | A | -758 |
| NW_005081596.1 | 1800347 | C/A | A | -755 |
| NW_005081596.1 | 1800648 | T/C | A | -454 |
| NW_005081596.1 | 1800745 | A/G | A | -357 |
| NW_005081596.1 | 1800769 | G/A | A | -333 |
| NW_005081596.1 | 1800798 | G/A | A | -304 |
| NW_005081596.1 | 1800878 | C/T | A | -224 |

**Table S8.** F and *p*-values for the omnibus test (2-way ANOVA) accompanying Fig.4B in the main text.

| | DT40 | | HEK-293 | | HeLa | |
|---|---|---|---|---|---|---|
| | *F* | *p* | *F* | *p* | *F* | *p* |
| CRE | 1.67 | 0.193 | 2.76 | 0.079 | 8.14 | < 0.001 |
| Allele | 22.92 | < 0.001 | 59.76 | < 0.001 | 1.13 | 0.29 |
| CRE x Allele | 1.74 | 0.18 | 2.76 | 0.079 | 7.1 | 0.001 |

**Table S9.** *P*-values for post-hoc comparisons (Student's pairwise t-test) accompanying Fig.4B in the

| CRE | DT40 | HEK-293 | HeLa |
|---|---|---|---|
| A | 0.019 | 0.002 | 0.002 |
| B | 0.016 | < 0.001 | 0.129 |
| C | 0.004 | 0.059 | 0.058 |

**Table S10.** CpGs in the *ESR1* CREs and Exon1A.

| Accession | Position | ZAL2/ZAL2$^{m}$ | Promoter | Annotation | Dist. from TSS |
|---|---|---|---|---|---|
| NW_005081596.1 | 1726299 | CG/TG | C | unshared_CpG_ZAL2 | -1531 |
| NW_005081596.1 | 1726431 | CG/CG | C | shared_CpG | -1399 |
| NW_005081596.1 | 1726483 | CG/CT | C | unshared_CpG_ZAL2 | -1347 |
| NW_005081596.1 | 1726858 | CG/CG | C | shared_CpG | -972 |
| NW_005081596.1 | 1727091 | CG/GG | C | unshared_CpG_ZAL2 | -739 |
| NW_005081596.1 | 1727328 | CG/CG | C | shared_CpG | -502 |
| NW_005081596.1 | 1748236 | CG/CG | B | shared_CpG | -1178 |
| NW_005081596.1 | 1748569 | CG/CG | B | shared_CpG | -845 |
| NW_005081596.1 | 1748650 | CT/CG | B | unshared_CpG_ZAL2$^{m}$ | -764 |
| NW_005081596.1 | 1748695 | CG/CG | B | shared_CpG | -719 |
| NW_005081596.1 | 1748948 | CA/CG | B | unshared_CpG_ZAL2$^{m}$ | -466 |
| NW_005081596.1 | 1749027 | CG/CG | B | shared_CpG | -387 |
| NW_005081596.1 | 1749148 | CG/CG | B | shared_CpG | -266 |
| NW_005081596.1 | 1749156 | CG/CG | B | shared_CpG | -258 |
| NW_005081596.1 | 1749208 | CG/CG | B | shared_CpG | -206 |
| NW_005081596.1 | 1749356 | CG/CG | B | shared_CpG | -58 |
| NW_005081596.1 | 1749414 | CG/CG | B | shared_CpG | 0 |
| NW_005081596.1 | 1799180 | CG/CG | A | shared_CpG | -1923 |
| NW_005081596.1 | 1799225 | CG/CG | A | shared_CpG | -1878 |
| NW_005081596.1 | 1799245 | CG/AG | A | unshared_CpG_ZAL2 | -1857 |
| NW_005081596.1 | 1799324 | CG/CG | A | shared_CpG | -1779 |
| NW_005081596.1 | 1799456 | CG/AG | A | unshared_CpG_ZAL2 | -1646 |
| NW_005081596.1 | 1799495 | CG/CG | A | shared_CpG | -1608 |
| NW_005081596.1 | 1799534 | CG/CG | A | shared_CpG | -1569 |
| NW_005081596.1 | 1799899 | CG/CG | A | shared_CpG | -1204 |
| NW_005081596.1 | 1800143 | CG/CG | A | shared_CpG | -960 |
| NW_005081596.1 | 1800387 | CG/CG | A | shared_CpG | -716 |
| NW_005081596.1 | 1800484 | CG/CG | A | shared_CpG | -619 |
| NW_005081596.1 | 1800768 | CG/CA | A | unshared_CpG_ZAL2 | -334 |
| NW_005081596.1 | 1800797 | CG/CA | A | unshared_CpG_ZAL2 | -305 |
| NW_005081596.1 | 1800856 | CG/CG | A | shared_CpG | -247 |
| NW_005081596.1 | 1800859 | CG/CG | A | shared_CpG | -244 |
| NW_005081596.1 | 1800884 | CG/CG | A | shared_CpG | -219 |
| NW_005081596.1 | 1800978 | CG/CG | A | shared_CpG | -125 |
| NW_005081596.1 | 1801023 | CG/CG | A | shared_CpG | -80 |
| NW_005081596.1 | 1801121 | CG/CG | Exon1A | shared_CpG | 18 |
| NW_005081596.1 | 1801127 | CG/CG | Exon1A | shared_CpG | 24 |
| NW_005081596.1 | 1801288 | CG/CG | Exon1A | shared_CpG | 185 |
| NW_005081596.1 | 1801295 | CG/CG | Exon1A | shared_CpG | 192 |
| NW_005081596.1 | 1801304 | CG/CG | Exon1A | shared_CpG | 201 |
| NW_005081596.1 | 1801369 | CG/CG | Exon1A | shared_CpG | 266 |
| NW_005081596.1 | 1801376 | CG/CA | Exon1A | unshared_CpG_ZAL2 | 274 |
| NW_005081596.1 | 1801393 | CG/CG | Exon1A | shared_CpG | 290 |
| NW_005081596.1 | 1801468 | CG/CG | Exon1A | shared_CpG | 365 |
| NW_005081596.1 | 1801477 | CG/CG | Exon1A | shared_CpG | 374 |
| NW_005081596.1 | 1801495 | CG/CG | Exon1A | shared_CpG | 392 |
| NW_005081596.1 | 1801509 | CG/CA | Exon1A | unshared_CpG_ZAL2 | 406 |
| NW_005081596.1 | 1801559 | CG/CG | Exon1A | shared_CpG | 456 |
| NW_005081596.1 | 1801574 | CG/CG | Exon1A | shared_CpG | 471 |

**Table S11.** *P*-values for post-hoc comparisons (estimated marginal means) controlling for individual accompanying Fig.4C in the main text.

| All sites | Effects of Allele | | |
|---|---|---|---|
| Allele | ZAL2 (TS) | ZAL2 (WS) | $ZAL2^{m}$ (WS) |
| ZAL2 (TS) | - | 0.933 | < 0.001 |
| ZAL2 (WS) | 0.933 | - | < 0.001 |
| $ZAL2^{m}$ (WS) | < 0.001 | < 0.001 | - |

| Shared Sites Only | Effects of Allele | | |
|---|---|---|---|
| Allele | ZAL2 (TS) | ZAL2 (WS) | $ZAL2^{m}$ (WS) |
| ZAL2 (TS) | - | 0. 903 | 0. 961 |
| ZAL2 (WS) | 0.903 | - | 0.984 |
| $ZAL2^{m}$ (WS) | 0.961 | 0.984 | - |

**Table S12.** Primers used to make *cis*-regulatory element (CRE) constructs for luciferase assays, and amplicons for next-generation bisulfite sequencing. Primers for luciferase assays included restriction cut sites for KpnI (F: GGTACC) or MluI (R: ACGCGT) on the 5' end of the primer. The bisulfite primers included Nextera transposase adapters on the 5' end of the primer (F: TCGTCGGCAGCGTCAGATGTGTATAAGAGACAG; R: GTCTCGTGGGCTCGGAGATGTGTATAAGAGACAG).

| Purpose | Target | Primer Name | Primer Sequence (5' ➔ 3') |
|---|---|---|---|
| Luciferase assay | A | ESR1A_F | TGCAGCTTAATAAGGGATGATGC |
| | | ESR1A_R | ACCAGAACCTATTCTGAGGCT |
| | B | ESR1B_F | GGTAAAGCAGGCCACTGTTAC |
| | | ESR1B_R | AGCAGGCTGTTGCCATTC |
| | C | ESR1C_F | GCCATATAAGCAAGAGGGC |
| | | ESR1C_R | CTCATTGGCAGTCACCAAG |
| Bisulfite sequencing | A | ESR1A_F_bs_89 | ATGTGTTAGGGTGATTGGTTTAAAT |
| | | ESR1A_R_bs_574 | CAATACTTTCAAAACTCCTACTAAAATAAC |
| | A | ESR1A_F_bs_96 | GGGTGATTGGTTTAAATTTGTAGTT |
| | | ESR1A_R_bs_527 | CAAAAATATTCTAAAACAATATATATTTTT |
| | A | ESR1A_F_bs_737 | GTTGGAGTTTATATATTAATGTATTTAGGG |
| | | ESR1A_R_bs_1179 | ATCATTTACATTTTATCAAAATTAAAAAAT |
| | A | ESR1A_F_bs_1269 | AGTGTTTTAGTAATTGTAAGTTGTTGTTGT |
| | | ESR1A_R_bs_1579 | CCTCCTATTTAATTCATTTTCATATTTACT |
| | A | ESR1A_F_bs_1656 | TTTGGTTGGATTTAGTGTTTTTTTT |
| | | ESR1A_R_bs_2116 | CAAAAAACATATCTACTTTTACTATCAACA |
| | B | ESR1B_F_bs_1828 | TTTTAATTTATTTGTTTAGTTTTTGATT |
| | | ESR1B_R_bs_2190 | CTAACTACTATAATAAATCACCAAATCTTC |
| | B | ESR1B_F_bs_1021 | AAAATTTTTGTTGGGGTATTTTGTA |
| | | ESR1B_R_bs_1370 | AACTAACTCACAAAACCTTCTAAATAAACA |
| | B | ESR1B_F_bs_1562 | TTGTTTTGGAAATAAGGTAATTTTGATA |
| | | ESR1B_R_bs_2001 | ACTATAAACACATCTCAAAAATTACTTTCA |
| | B | ESR1B_F_bs_2055 | AAATTATTGGAAATTTTAGTTTTAAGA |
| | | ESR1B_R_bs_2494 | AACAAACTATTACCATTCCCTCTAC |
| | C | ESR1C_R_bs_424 | GAAGGAAGGGTTTGTTTATTATGATT |
| | | ESR1C_F_bs_837 | TCCATCTCCTCAAAATAATATCTAATATCT |
| | C | ESR1C_R_bs_915 | AGGAGGAATAGATTAATGTAAGGAATTAAA |
| | | ESR1C_F_bs_1313 | TCAACACCTACATTACAAAATATAAATAAA |
| | C | ESR1C_F_bs_1226 | AAGTTAGTAGATAGTAGAGAAGTTAGAAGA |
| | | ESR1C_R_bs_1622 | AAATAATTTAACACAATTTCTTTCCC |
| | Exon 1A | ESR1E1_F_bs_129 | AAGGTATTGAATTGGAGATATTGAGTAG |
| | | ESR1E1_R_bs_499 | AACTAAAAAAACAACTTCCCTCATC |

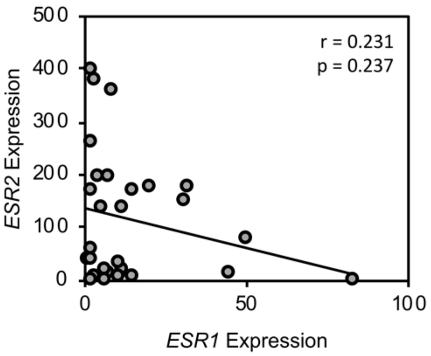


**Figure S1.** The relationship between *ESR1* and *ESR2* expression, as measured by qPCR, in nucleus taeniae (TnA) of birds in the knockdown experiment (see Fig. 2). Only birds receiving scrambled control oligonucleotides or for whom the antisense infusions missed TnA are included here.

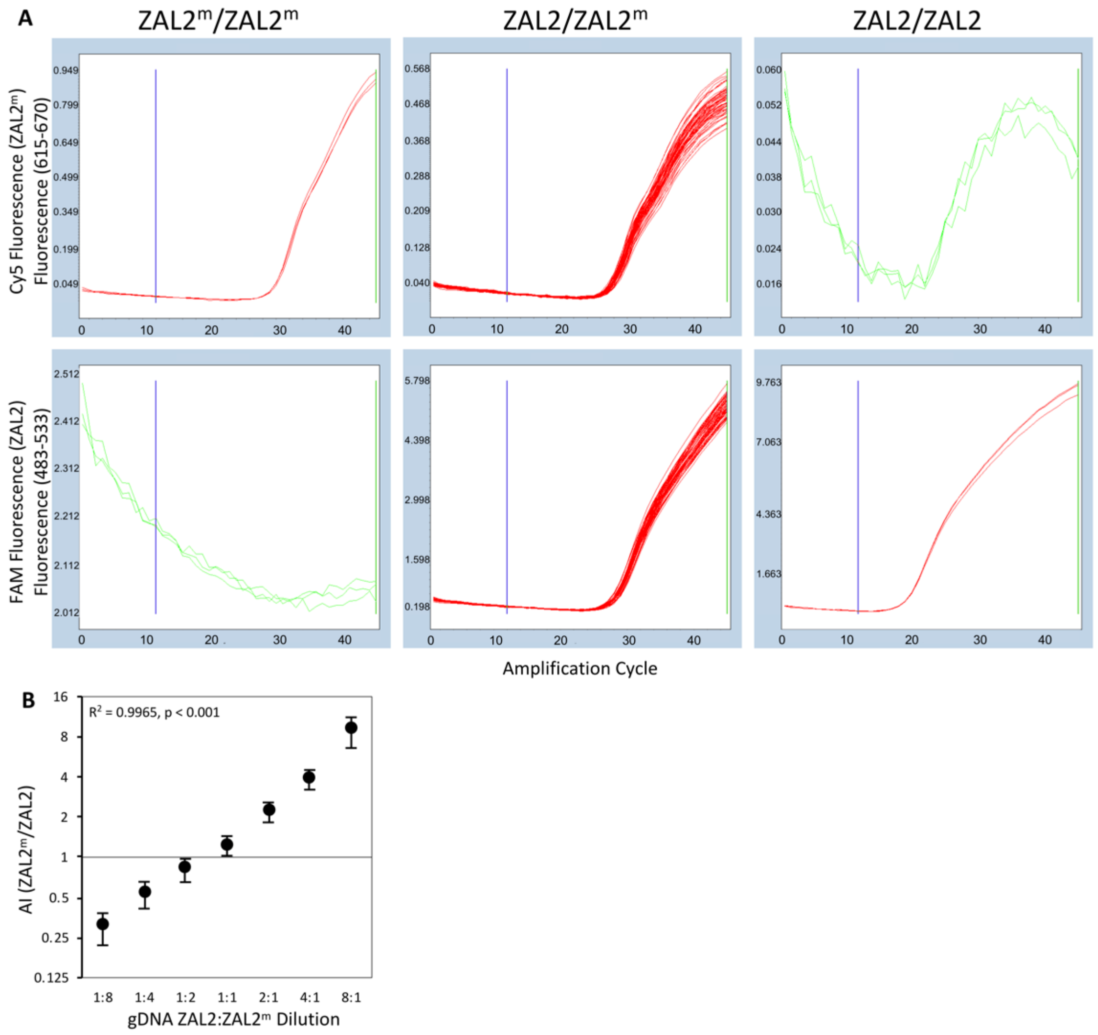


**Figure S2**. Validation of the AI assay. (A) Allele-specific qPCR amplification curves for ZAL2 and ZAL2$^{m}$ genomic DNA, demonstrating that the probes selectively amplify the targeted allele (ZAL2$^{m}$/ZAL2$^{m}$, n = 1; ZAL2/ZAL2$^{m}$ n = 12, ZAL2/ZAL2 n = 1; run in triplicate). Note the difference in the Y-axis scale in the top right and bottom left (green) – no amplification curve was observed. (B) An example of an AI standard curve of dilutions from 1:8 to 8:1 of ZAL2: ZAL2$^{m}$ genomic DNA. Mean + sem, n = 3 per dilution ($y = 0.1644e0.5403^{x}$).

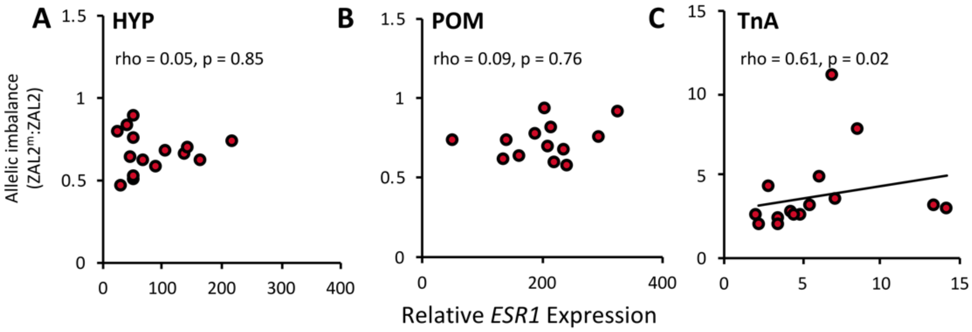


**Figure S3**. Spearman rank correlations show that allelic imbalance of *ESR1* does not predict *ESR1* expression in (A) hypothalamus (HYP) or (B) the medial preoptic area (POM) but it does in (C) nucleus taeniae (TnA). *ESR1* expression for these animals, as measured by qPCR, was published by Grogan et al.[15]

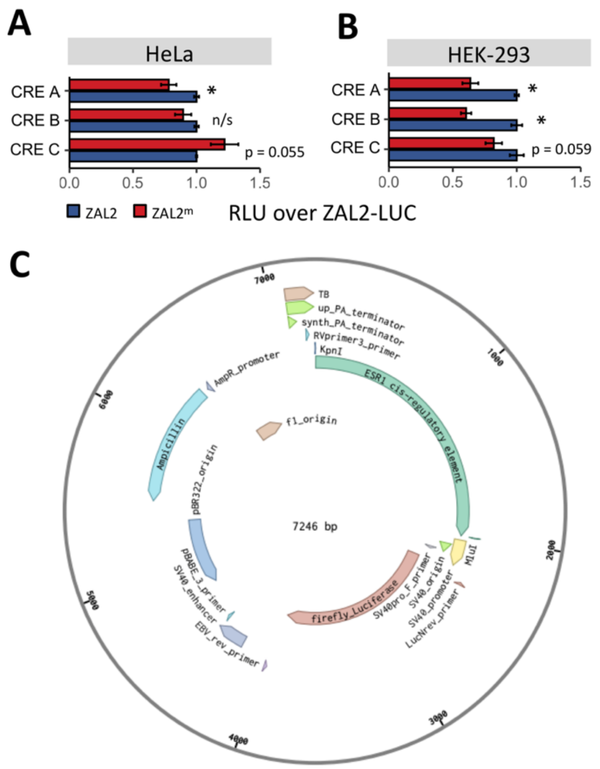


**Figure S4**. Luciferase assays. (A,B) Effect of *cis*-regulatory variation in *ESR1* on luciferase activity *in vitro* in HeLa (A) and HEK-293 (B) cells. The activity, in relative light units (RLU), of the ZAL2$^{m}$-luciferase (LUC) (red) and ZAL2-LUC (blue) constructs are shown normalized to activity of the ZAL2-LUC construct. * $p < 0.05$; see Table S10 and S11 for all p values. The direction of the effect of allele was opposite in HEK cells, compared with HeLa cells (A) and DT40 cells (Fig. 4B, main text). This result is consistent with variation in local availability of transcription factors in lines of cultured cells[16]. The local complement of transcription factors in each cell line is expected to interact differently with the ZAL2 and ZAL2$^{m}$ alleles (see Fig. S5), which could change the direction of the overall effect of allele. (C) The *ESR1 cis*-regulatory elements (CREs) were cloned into the pGL3-control vector (Genbank E1741) at KpnI and MluI cut sites upstream of the firefly_luciferase gene.

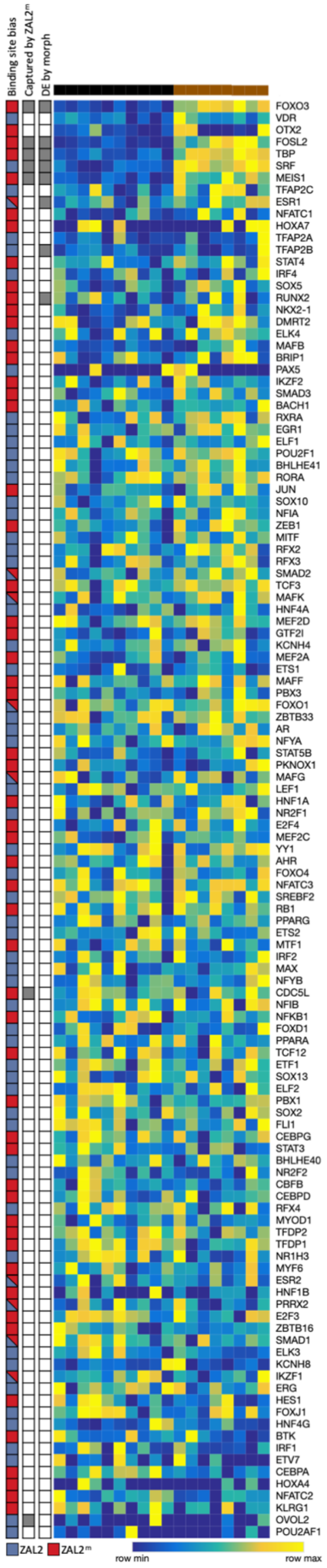


**Figure S5**. Expression of 120 transcription factors (TFs) with binding sites disrupted by ZAL2/ZAL2$^{m}$ fixed SNPs (see Fig. 4A). Gene expression is shown from RNA-seq data[8,10] normalized using "estimateSizeFactors" in DeSeq2 in WS (black) and TS (tan) males. RNA was extracted from nucleus taeniae (TnA). Each row represents a gene and each column represents an individual. Genes are ordered according to fold difference between WS and TS birds. TFs that are differentially expressed by morph (Benjamini-Hochberg correction at FDR = 0.1), or captured by ZAL2$^{m}$, are marked with a light grey box to the left. On the far left, TFs are color-coded red or blue according to allelic bias, in other words according to which allele contains one or more binding sites predicted to have higher affinity for that particular TF than the corresponding site(s) on the other allele. Eight TFs are coded as both blue and red because there are multiple disrupted binding sites; for half of the sites, the ZAL2 allele is predicted to have higher affinity for transcription factors whereas for the rest, ZAL2$^{m}$ is predicted to have higher affinity.

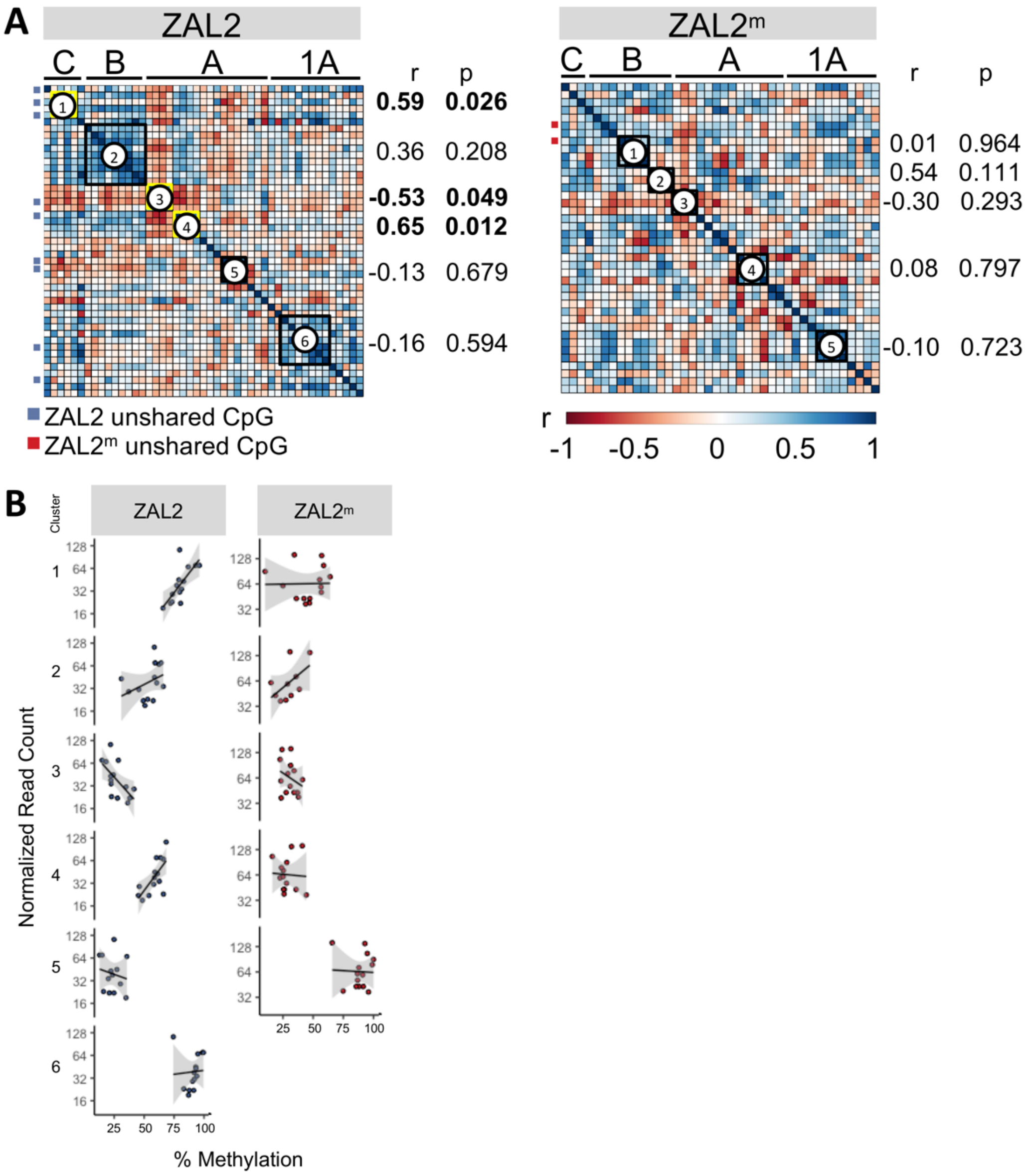


**Figure S6**. Allele-specific methylation predicts allele-specific expression. (A) Correlation matrices using data from WS birds are shown (see Fig. 4). Similarly methylated clusters of sites are enclosed in boxes and numbered (see Fig. 4E in the main text for the figure with clusters unobscured). The clusters were used to test the association between allele-specific methylation and allele-specific expression in the same animals. Clusters that significantly predicted allele-specific expression are outlined in yellow, clusters that did not are outlined in black. Unshared sites are marked by blue (ZAL2) or red (ZAL2$^{m}$) boxes to the left of each matrix to indicate which allele contains the site. (B) Scatterplots show the relationships between allele-specific methylation and allele-specific expression (from RNA-seq data; see Methods) for the clusters shown in (A). * $p < 0.05$.
